# UK-Biobank Whole Exome Sequence Binary Phenome Analysis with Robust Region-based Rare Variant Test

**DOI:** 10.1101/697912

**Authors:** Zhangchen Zhao, Wenjian Bi, Wei Zhou, Peter VandeHaar, Lars G. Fritsche, Seunggeun Lee

## Abstract

In biobank data analysis, most binary phenotypes have unbalanced case-control ratios, which can cause inflation of type I error rates. Recently, a saddlepoint approximation (SPA) based single variant test has been developed to provide an accurate and scalable method to test for associations of such phenotypes. For gene- or region-based multiple variant tests, a few methods exist which adjust for unbalanced case-control ratios; however, these methods are either less accurate when case-control ratios are extremely unbalanced or not scalable for large data analyses. To address these problems, we propose SKAT/SKAT-O type region-based tests, where the single-variant score statistic is calibrated based on SPA and Efficient Resampling (ER). Through simulation studies, we show that the proposed method provides well-calibrated p-values. In contrast, the unadjusted approach has greatly inflated type I error rates (90 times of exome-wide *α* =2.5×10^-6^) when the case-control ratio is 1:99. Additionally, the proposed method has similar computation time as the unadjusted approaches and is scalable for large sample data. Our UK Biobank whole exome sequence data analysis of 45,596 unrelated European samples and 791 PheCode phenotypes identified 10 rare variant associations with p-value < 10^-7^, including the associations between *JAK2* and myeloproliferative disease, *TNC* and large cell lymphoma and *F11* and congenital coagulation defects. All analysis summary results are publicly available through a web-based visual server.

## Introduction

With the decreased cost of sequencing, big biobanks have started to whole exome or whole genome sequence large number of participants to identify the role of rare variants to complex diseases^1-3^. By combining rich phenotypic information in electronic health record (EHR)^4^, these sequence data will illuminate the phenome-wide association patterns of rare variants. Since most of diseases and symptoms have low prevalence, the binary phenotypes in biobanks generally have unbalanced case-control ratios (1:10 or 1:100, for example)^5^. For example, in the UK Biobank data, nearly 99% of PheCode-based binary phenotypes have case-control ratios less than 1:10 ^6^. Substantial challenges are posed when analyzing the associations between rare variants and unbalanced phenotypes.

Since single-variant tests are underpowered to identify disease associated rare variants^7^, gene- or region-based multiple variant tests, including burden test^8,9^, SKAT^10^, and SKAT-O^11^, are commonly used to identify rare variant associations. To evaluate the association signals in multiple variants, these methods aggregate single variant score statistics. However, as shown in our simulation studies and elsewhere^12-14^, these methods suffer from the inflation of type I error rates when case-control ratios are unbalanced. For single variant tests, saddlepoint approximation (SPA) based approach has been developed and provides accurate p-values under such a case-control imbalance^5,15^. Although a few methods exist which adjust for unbalanced case-control ratios for gene- or region-based tests, including moment-based adjustment (MA)^16^ and efficient resampling (ER)^16^, these methods are not scalable or accurate for biobank data. When the case-control ratio is extremely unbalanced, MA can still have inflated type I error rates. ER is computationally expensive when minor allele counts (MAC) are moderate or large. To address these problems, we propose a robust region-based test that adjusts single variant score statistics using SPA and ER, and aggregate the adjusted statistics. The SPA and ER help to precisely calculate the reference distribution of the single variant score statistics, thereby properly controlling for the type I error rates. The computation cost of the proposed approach is comparable to unadjusted tests, and hence can be applied to large biobank data. Using extensive simulation studies, we demonstrate that our robust burden, SKAT, and SKAT-O tests have proper type I error rates even when the case-control ratio is 1:99 and exhibit larger power compared to the unadjusted burden, SKAT, and SKAT-O test. In addition, the method can be applicable not only rare variant tests but also the joint association test of common and rare variants.

The UK Biobank resource^2^ was extended with the first tranche of whole exome sequencing (WES) data for 49,960 participants^1^. We performed robust gene-based rare-variant tests of 45,596 unrelated European samples on 791 phenotypes with at least 50 cases and identified 10 rare variant associations with p-value < 10^-7^, including the associations between *JAK2* and myeloproliferative disease, *NAGS* and cervical intraepithelial neoplasia, *TNC* and large cell lymphoma. These results shed light on the discoveries we can make with full 500,000 WES samples, which will be available in near future.

## Results

The proposed approach calibrates single variant score statistics for region-based tests. The calibration is performed using SPA and ER. SPA is an asymptotic-based approximation to the true distribution of score statistics by approximating the inversion of the cumulant generating function^17,18^. It has a faster convergence rate than using normal distribution^5^, but when the minor allele count is too low (ex. MAC < 10), the method cannot work properly. ER is a resampling-based approach and provides an exact p-value when MAC is low^16^. However, as MAC increases, the computation cost increases rapidly. The proposed approach combines these two methods: when the variant MAC < 10, ER is used to calculate the p-values of single variant score statistics, and when MAC >=10, SPA is used. The p-values are used to calibrate the variance estimates, and then the gene- and region-based p-value is calculated with the updated variance. The details can be found in Methods below.

### Type I Error and Power Simulation Results

We generated 10^7^ datasets to compare type I error rates of the proposed approaches (Robust burden, SKAT and SKAT-O), unadjusted approaches (burden, SKAT and SKAT-O) and a hybrid approach for SKAT-O^16^. The hybrid approach applies several adjustment methods based on MAC. Table 1 shows that the unadjusted approaches had substantial inflation of type I error rates when the case-control ratio was unbalanced. In contrast, the robust SKAT controlled type I error rates much better and had only a slight inflation when the case-control ratio was 1:99. Interestingly, the existing hybrid approach showed substantially inflated type I error rates when case-control ratios were extremely unbalanced (case-control ratio=1:49 and 1:99). This may be due to the fact that the MAC-based method selection rule in the hybrid approach do not perform well under extremely unbalanced case-control ratios. When the case-control ratios are more extreme than 1:99, the robust SKAT and SKAT-O showed some inflation of type I error rates (Supplementary Table 1). Overall, the type I error simulation results confirmed that the proposed robust approaches provide substantially improved type I error rates compared to the unadjusted and the existing hybrid approaches.

**Table 1.** Type I error rates of unadjusted and robust versions of SKAT, burden and SKAT-O and hybrid method when testing rare variants with dichotomous traits at *α* = 10^−2^, 10^−4^ and 2.5 × 10^−6^. The sample size was 50,000 and 10^7^ datasets were generated.

| $\alpha$ | Case:<br>control | SKAT | Robust<br>SKAT | Burden | Robust<br>burden | SKAT-O | Robust<br>SKAT-O | Hybrid<br>SKAT-O |
| --- | --- | --- | --- | --- | --- | --- | --- | --- |
| $10^{-2}$ | 1:1 | 0.99 | 0.99 | 1.00 | 1.00 | 1.11 | 1.11 | 1.09 |
|  | 1:9 | 1.01 | 1.01 | 0.99 | 1.00 | 1.13 | 1.13 | 1.09 |
|  | 1:49 | 1.44 | 1.22 | 1.02 | 0.95 | 1.44 | 1.23 | 1.27 |
|  | 1:99 | 1.92 | 1.41 | 1.07 | 0.91 | 1.82 | 1.33 | 1.53 |
| $10^{-4}$ | 1:1 | 0.99 | 1.03 | 1.02 | 1.00 | 1.27 | 1.32 | 1.27 |
|  | 1:9 | 1.39 | 1.14 | 1.12 | 0.99 | 1.65 | 1.40 | 1.52 |
|  | 1:49 | 6.31 | 1.65 | 2.43 | 0.97 | 6.16 | 1.79 | 4.54 |
|  | 1:99 | 13.48 | 2.13 | 3.95 | 1.02 | 12.77 | 2.17 | 8.89 |
| $2.5 \times 10^{-6}$ | 1:1 | 1.24 | 1.54 | 1.11 | 1.03 | 1.38 | 1.38 | 1.40 |
|  | 1:9 | 2.47 | 1.45 | 1.29 | 0.77 | 2.51 | 1.49 | 2.23 |
|  | 1:49 | 28.27 | 1.91 | 6.88 | 1.06 | 23.70 | 1.98 | 16.69 |
|  | 1:99 | 89.53 | 1.81 | 16.34 | 0.90 | 71.32 | 1.60 | 42.59 |

Figure 1 shows the empirical powers of the hybrid, unadjusted and robust version of SKAT-O methods, considered at type I error simulations. The empirical powers of unadjusted and robust versions of SKAT and burden can be found in Supplementary Figure 1. Since unadjusted and hybrid methods had severely inflated type I error rates, for the fair comparison, we used the empirical significance level estimated from type I error simulation studies. Assuming that the type I error rates could be properly controlled for all methods, robust SKAT-O had similar power as unadjusted SKAT-O in balanced and moderately unbalanced case-control ratios (1:1 and 1:9) and was more powerful than unadjusted SKAT-O in extremely unbalanced ratios (1:49 and 1:99). Robust burden tests had the same power as unadjusted burden tests across all four case-control ratios. Robust SKAT had similar power with unadjusted SKAT in balanced ratios and was more powerful than unadjusted SKAT in unbalanced ratios. If the number of cases was fixed, more controls (1:49 and 1:99) increased power greatly compared to case-control ratio 1:1 for all three robust methods (Supplementary Figure 2). In addition, we found that 1:99 had slightly more power than 1:49, where we could infer that 1:99 is sufficient to achieve the maximum power and more controls can hardly increase the power.

**Figure 1.**
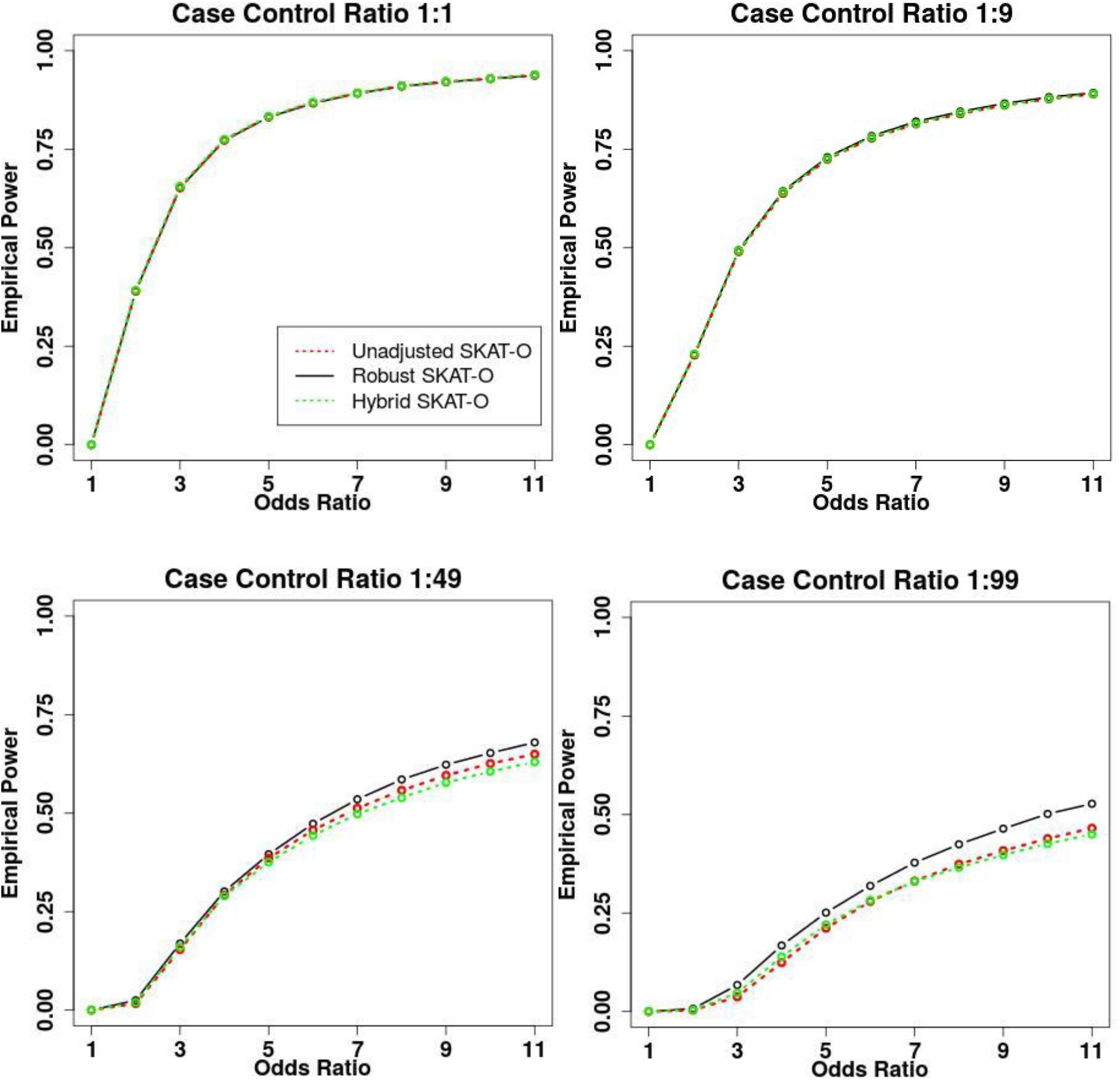
Empirical power estimates for the unadjusted and robust versions of SKAT-O and hybrid method where 30% of variants were causal variants and all causal variants were deleterious. The sample size was 50,000 and 10,000 datasets were generated. The X-axis represents the genetic effect odds ratio and the Y-axis represents the empirical power. All causal variants had the same odds ratios.

**Figure 2.**
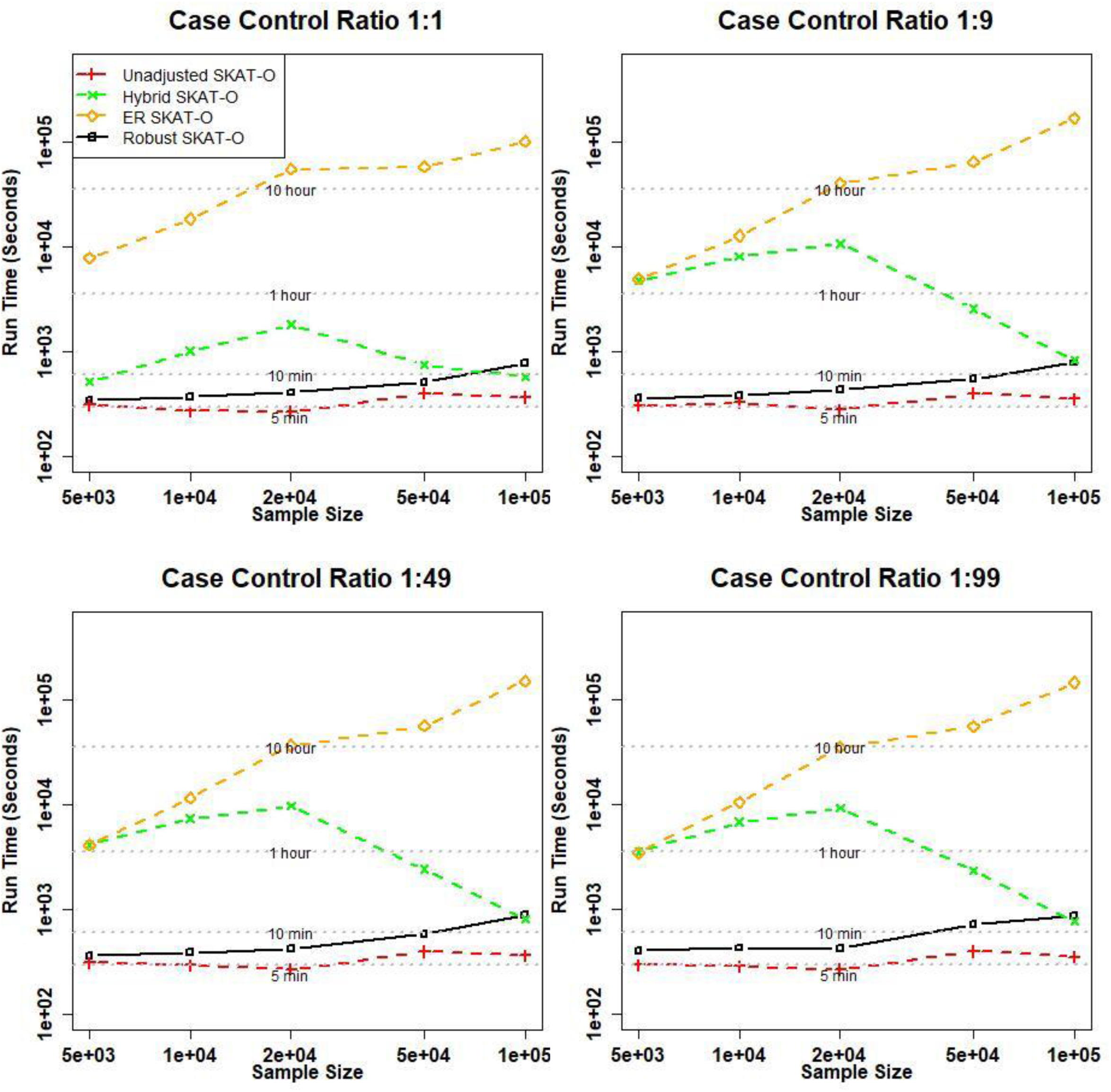
Comparison of computation time of unadjusted, hybrid, ER and robust approaches for SKAT-O. The rare-variant region-based tests were performed on randomly selected 1 kb regions of 1,000 resamples. The X-axis represents the sample size and the Y-axis represents the run time of 1,000 resamples.

In summary, the robust methods had similar or more power than the unadjusted methods in all scenarios. Among the three robust methods, robust SKAT-O generally performed better than robust SKAT and robust burden tests since robust SKAT-O combined the two tests (Supplementary Figure 3).

**Figure 3.**
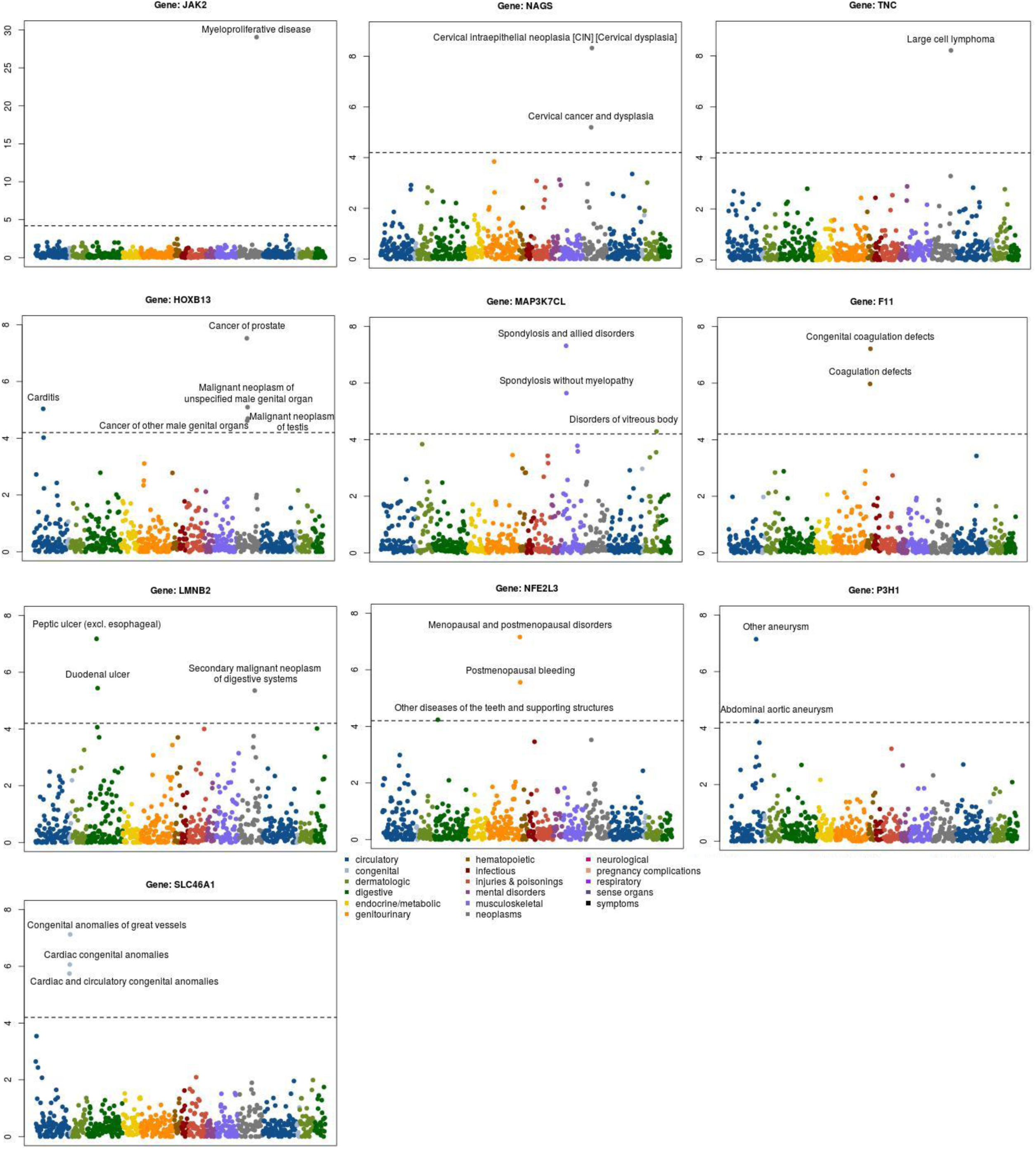
PheWAS plots of 10 rare variant associations with p-value< 10^-7^. The X-axis represents 791 binary traits and the Y-axis represents the negative log10 p-values. The dashed line represents the cutoff of 0.05/791=6.32×10^-5^.

### Comparison of computational times

To compare the computation times, we generated 1,000 datasets (Figure 2). Since SKAT-O combines the burden and SKAT tests, we only considered the SKAT-O test. As the sample sizes increased, the computation time of ER increased and required ∼16.1 CPU hours for analyzing one gene for 50,000 individuals. In contrast, unadjusted methods required 140x less computation time (∼6.7 min) and the computation times barely changed by sample size (5,000-100,000 individuals). Our robust method performed similarly as unadjusted SKAT-O (∼8.5 min). Since the hybrid approach selects its methods based on MAC and case-control ratios, the computation cost of the hybrid approach is not determined by the sample size. Overall, the hybrid approach was slower than the proposed method.

The computation time for analyzing UK-Biobank data of 791 binary phenotypes with robust SKAT-O was 453 CPU days, i.e. ∼13.7 CPU hours per one phenotype.

### Analysis of whole exome sequencing (WES) data in the UK Biobank

We applied six methods (unadjusted and robust versions of SKAT, burden and SKAT-O tests) to the analysis of WES data in the UK Biobank. We restricted our analysis to the rare nonsynonymous and splicing variants with minor allele frequencies (MAFs) < 0.01 in exon regions. A total of 18,360 genes were analyzed based on 45,596 independent European samples across 791 binary phenotypes with at least 50 cases. For phenotypes with case-control ratios more extreme than 1:99, we identified the ancestry-matched control samples to make case-control ratios 1:99 (See Method).

With the cutoff of *α* = 2.5 × 10^−6^, unadjusted SKAT-O detected 77,941 significant genes, most of them would be false positives, while our robust methods detected 102 significant genes for SKAT, 40 for the burden test and 117 for SKAT-O. Since we were testing many phenotypes, the usual exome-based cutoff of 2.5 × 10^−6^ can produce spurious associations. Following Hout et al^1^, we used a more stringent level *α* =10^-7^ and identified that 10 gene-phenotype pairs had robust SKAT-O p-values smaller than 10^-7^ (Table 2). Among them, rare variant associations between *JAK2* and myeloproliferative disease (number of cases=94)^19^, and *HOXB13* and prostate cancers (number of cases=741)^20^ have been previously reported, which demonstrates that our analysis can replicate known signals, even when the number of case samples is very small. Among 10 phenotype-gene pairs, only 2 had a single SNP p-value < 5 × 10^−8^, indicating that gene/region-based approaches are more powerful than single variant analyses. For each gene, the top 3 smallest p-value variants were reported in Supplementary Table 4 and single variant p-values were presented in Supplementary Figure 4. QQ plots for those 10 phenotypes show that unadjusted SKAT-O had greatly inflated type I error rates, but our robust approach provided relatively well calibrated results (Supplementary Figure 5).

**Table 2.** Significant Gene-Phenotype Associations Based on UK Biobank WES Data. Lowest P SNP means the lowest p-value of all single variants contained in the gene-phenotype association. Conditional P-value (SKAT-O) means the robust SKAT-O p-value after conditioning on the most significant nearby variants (± 100 Kbp up and down stream). P-value of the most significant nearby variant was from SAIGE single variant analysis results^15^ of the UK-Biobank imputed datasets of 400,000 British samples.

| Phenotype (PheCode) | Gene name | Case: control | The number of snps | Total minor allele counts for cases | Total minor allele counts for controls | Robust SKAT-O P-values | Lowest P SNP | Conditional P-value (SKAT-O) | P-value of the most significant nearby variant |
| --- | --- | --- | --- | --- | --- | --- | --- | --- | --- |
| Myeloproliferative disease (200) | *JAK2* | 94: 9306 | 68 | 26 | 435 | $8.92 \times 10^{-30}$ | $4.39 \times 10^{-36}$ | $6.69 \times 10^{-32}$ | $2.30 \times 10^{-17}$ |
| Cervical intraepithelial neoplasia [CIN] [Cervical dysplasia] (180.3) | *NAGS* | 309:21875 | 81 | 21 | 340 | $4.71 \times 10^{-9}$ | $4.37 \times 10^{-4}$ | $1.33 \times 10^{-8}$ | $2.12 \times 10^{-3}$ |
| Large cell lymphoma (202.24) | *TNC* | 56:5544 | 135 | 14 | 491 | $6.10 \times 10^{-9}$ | $1.02 \times 10^{-5}$ | $2.99 \times 10^{-9}$ | $7.89 \times 10^{-4}$ |
| Cancer of prostate (185) | *HOXB13* | 741:18940 | 37 | 18 | 154 | $3.00 \times 10^{-8}$ | $5.24 \times 10^{-8}$ | $5.67 \times 10^{-8}$ | $1.12 \times 10^{-4}$ |
| Spondylosis and allied disorders (721) | *MAP3K7CL* | 849:43787 | 58 | 14 | 153 | $4.85 \times 10^{-8}$ | $2.11 \times 10^{-7}$ | $7.59 \times 10^{-9}$ | $3.46 \times 10^{-3}$ |
| Congenital coagulation defects (286.1) | *F11* | 76:7524 | 39 | 8 | 85 | $6.13 \times 10^{-8}$ | $4.52 \times 10^{-5}$ | $3.83 \times 10^{-8}$ | $4.21 \times 10^{-3}$ |
| Peptic ulcer (excl. esophageal) (531) | *LMNB2* | 773:44818 | 171 | 24 | 501 | $6.65 \times 10^{-8}$ | $3.83 \times 10^{-6}$ | $6.27 \times 10^{-8}$ | $4.17 \times 10^{-1}$ |
| Menopausal and postmenopausal disorders (627) | *NFE2L3* | 1345:21226 | 172 | 144 | 1369 | $6.93 \times 10^{-8}$ | $2.72 \times 10^{-5}$ | $1.85 \times 10^{-7}$ | $2.14 \times 10^{-4}$ |
| Other aneurysm (442) | *P3H1* | 164:16236 | 112 | 17 | 498 | $7.13 \times 10^{-8}$ | $1.71 \times 10^{-5}$ | $2.26 \times 10^{-7}$ | $5.12 \times 10^{-4}$ |
| Congenital anomalies of great vessels (747.13) | *SLC46A1* | 134:13266 | 29 | 11 | 256 | $7.50 \times 10^{-8}$ | $1.86 \times 10^{-8}$ | $1.97 \times 10^{-8}$ | $3.71 \times 10^{-4}$ |

Among other genes, *NAGS* causes N-acetylglutamate synthase deficiency, an autosomal recessive disorder of the urea cycle^21^. In our data, *NAGS* was significantly associated with cervical intraepithelial neoplasia (p-value=4.71×10^-9^). PheWAS plot (Figure 3) also shows an association signal between *NAGS* and cervical cancer (p-value= 6.37×10^-6^). These findings are supported by recent literature which has shown that urea cycle dysregulation is related to cancer^22^. The *TNC* gene encodes Tenascin-C, an extracellular matrix protein with a spatially and temporally restricted tissue distribution. *TNC* has been associated with large cell lymphoma (p-value=6.10×10^-9^) consistent with the finding that Tenascin-C is highly expressed in various tumors including T-Cell non-Hodgkin lymphomas^23^ and associated with invasive front of tumors^24^. Although *TNC* is a well-known cancer maker, to the best of our knowledge, this is the first report of the role of rare variants in *TNC* in lymphoma. *F11*, also known as Coagulation Factor XI, was observed as associated with congenital coagulation defects (p-value=6.13×10^-8^), which is consistent with the fact that Factor XI participates in blood coagulation as a catalyst in the conversion of factor IX to factor IXa in the presence of calcium ions^25^.

We carried out conditional analysis to evaluate whether the rare variant association signals were independent of the nearby variant association signals (± 100 Kbp up and down stream) (Table 3). To identify most significant nearby variants, we used SAIGE single variant analysis results of the UK-Biobank imputed datasets of 400,000 British samples^15^. All ten associations remained significant after the conditional analysis.

We have generated summary statistics for all gene-phenotype association results using our robust approach and made them available in a PheWEB like visual server (See Code and data availability).

## Discussion

In this paper, we present a robust approach that can address case-control imbalance in region-based rare variant tests. The proposed approach uses recently developed ER and SPA to calibrate the variance of single variant score statistics to accurately calculate region-based p-values. Computation cost of the proposed approach is similar to the unadjusted approach, which makes it scalable for large analysis. Simulation studies showed that unadjusted methods suffer severe inflation of type I error rate in unbalanced case-control ratios while robust methods can successfully address it. The UK-Biobank exome data analysis shows that the method provides calibrated p-values and contribute to identifying true association signals.

The proposed robust methods combine SPA and ER to recalibrate variances of single score statistics. SPA can be thought as higher order asymptotic approach with error bound *O*(*n*^−3/2^) ^5^, where *n* is the sample size, which is much smaller than the error bound of normal approximation, *O*(*n*^−1/2^). But SPA is still asymptotic-based and cannot perform well when MAC is small. Since ER is a resampling-based approach and can calculate the exact p-value when MAC is small, it can complement SPA.

Our UK Biobank WES data analysis of 45,596 European samples have identified 10 rare variant associations with p-value < 10^-7^, including the replication of two known signals. Currently UK-Biobank is carrying out whole exome sequencing for 500,000 individuals. Our analysis shows the early snapshot of the discoveries that can be made with full UK-Biobank samples. All the UK-Biobank analysis summary statistics are publicly available, which can be a useful community resource to show detailed results of the UK-Biobank. For example, researchers could utilize it for meta-analysis to combine samples with different studies. It can also be used to validate novel signals from other studies.

There are several limitations in the proposed method. Currently, the robust methods require all individuals are unrelated. When there are related individuals, generalized linear mixed model (GLMM) based approaches^15,26^ should be used to incorporate the relatedness. Recently Wei et al developed scalable GLMM for gene-based tests that can handle the full size of UK-Biobank data of 500,000 samples^27^. In future, we will apply the robust approach to gene-based GLMM. Second, when the case-control ratios are more extreme than case:control=1:99, the method suffered type I error inflation. Because of this, our UK-Biobank exome analysis used the matching scheme in which if the case control ratios are more extreme than 1:99, we use the matching to reduce the number of controls.

In summary, we have proposed a robust region-based method and showed that the method can accurately analyze UK-Biobank exome data. With the continuous decrease of sequencing cost and growing effort to build large biobanks and cohorts^28^, rare variants association analysis will be increasingly applied to binary phenome. Our method will provide accurate results for binary phenome analysis and contribute to finding the role of rare variants to complex diseases.

## Methods

### Gene/region-based rare variant tests for binary traits

Assume *n* individuals are sequenced in a region, which has *m* rare variants. For the *i*-th individual, let *y*_*i*_ denote a binary phenotype, *G*_*i*_ = (*g*_*i*1_, *g*_*i*2_, …, *g*_*im*_)′ the hard call genotypes (*g*_*ij*_ = 0,1,2) or dosage values of the *m* genetic variants in the target gene or region, and *X*_*i*_ = (*X*_*i*1_, *X*_*i*2_, …, *X*_*is*_)′ the covariates, including the intercept. To model the binary outcome, the following logistic regression model can be used:

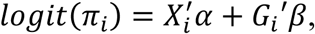

where *π*_*i*_ is the disease probability for the *i*-th individual, *α* is an *s* × 1 vector of regression coefficients of covariates, and *β* is an *m* × 1 vector of regression coefficients of genetic variants. Suppose 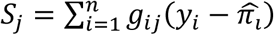 is the score statistic for the variant *j*, where 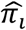 is the estimated disease probability under the null hypothesis of no association (i.e. *β* = 0). Burden and SKAT test statistics can be written as

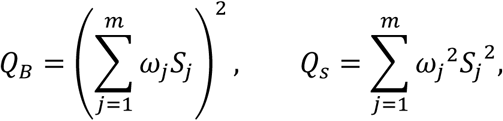

where *w*_*j*_ is the weight for each variant.^10^ In the simulation and real data analysis, we used the beta(1,25) weight, which upweight rarer variants^10^. The SKAT-O method combines the burden test and SKAT with the following framework:

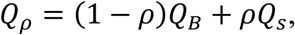

where *ρ* is a tuning parameter with range [0,1]. Since the optimal *ρ* is unknown, SKAT-O applies the minimum p-values over a grid of *ρ* as a test statistic.

Under the null hypothesis, *S* = (*S*_1_, …, *S*_*m*_)^*T*^ asymptotically follows the multivariate normal distribution, 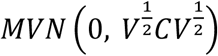, where *C* is the correlation matrix among *m* variants and *V* is a diagonal matrix where the diagonal elements are the asymptotic variances of *S*. In the presence of a case-control imbalance, however, the distribution of score statistics is skewed, which causes the inflation of type I error rates. To address this problem, we will utilize SPA and ER to adjust the variance matrix *V*.

### Saddle Point Approximation (SPA) and Efficient Resampling (ER)

SPA is a statistical method to calculate the distribution function using the cumulant generating function (CGF). Since it utilizes all the cumulants, SPA is more accurate than using normal approximation, which only uses the first two cumulants (mean and variance). Drawing on the work of Dey et al^5^, suppose *K*_*j*_(*t*) is the CGF of the score statistic *S*_*j*_, which can be derived based on the fact that *Y*_*i*_∼Bernoulli(*π*_*i*_) under the null. Then, the distribution function of the score statistic *S*_*j*_ can be approximated by

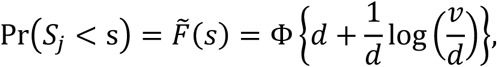

where 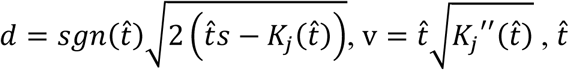, is the solution to the equation 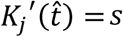, and φ is the distribution function of the standard normal distribution^5^.

Although SPA performs better than normal approximation, since it is still an asymptotic-based approach, SPA can result in inaccurate p-values when MAC is very low. To address this issue, we use ER for low MAC variants. ER is a resampling method that resamples the case–control status of individuals with a minor allele at a given variant and disease risk *π*_*i*_ instead of permuting case–control status across all individuals. This is because only individuals with minor alleles contribute to the score statistics *S*. Since ER is resampling-based, it can provide an accurate p-value for a very rare variant. When MAC is low (ex. MAC ≤ 10), ER can rapidly calculate the exact p-value by numerating all possible configurations of case-control statuses. The detailed derivations of ER can be found in Lee et al^16^.

### Robust SKAT, Robust burden test and Robust SKAT-O

For each variant *j*, when the score statistic *S*_*j*_ lies within 2 standard deviations of the mean, the normal approximation generally performs well^5^. Otherwise, due to the skewed distribution, the normal approximation causes inflated type I error rates. Hence, when *S*_*j*_ is out of 2 standard deviations of the mean, we apply SPA (when MAC > 10) or ER (when MAC ≤ 10) to calculate the p-value 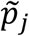, which will be used to calibrate the variance of *S*_*j*_.

Let 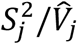 be a square-standardized test statistic in which 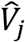 is the estimated variance of 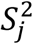. When *S*_*j*_ follows the normal distribution, 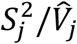 follows the chi-square distribution with one degree of freedom. We adjust the variance as the p-value is the same as 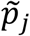, in which the adjusted variance is

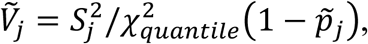

where 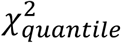 is the quantile function of the chi-square distribution with one degree of freedom. Note that if *S*_*j*_ lies within 2 standard deviations of the mean, 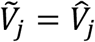. Suppose 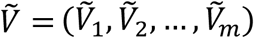, then the p-value of the region can be calculated based on the assumption that

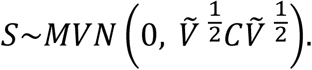

The adjustments above overcome the inflated type I error rates for common variants, but are insufficient to address the inflation issue for rare variants (4.87 times of exome-wide alpha=2.5×10^-6^ when the case-control ratio is 1:99. Details can be found in Supplementary Table 1). We apply additional adjustment by using the fact that burden test can be presented as a single marker test with collapsed variants, and SPA performs very well for single marker test. From the above equation, the variance estimate of the burden test is 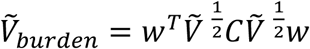, where *w* = (*w*_1_, …, *w*_*m*_)^*T*^ is an *m* × 1 vector of the weight. Suppose 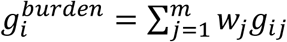, and then the burden test statistic (i.e. *Q*_*B*_) is identical to 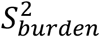, where 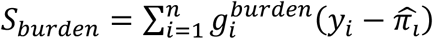, and the p-value 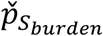 of *S*_*burden*_ can be calculated from SPA. Using the similar approximation in the above, we estimate the variance *S*_*burden*_ as 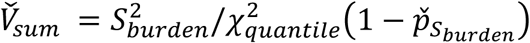. Suppose 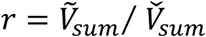. In order to control type I error inflation, we suggest utilizing a more conservative variance. Let 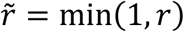, then

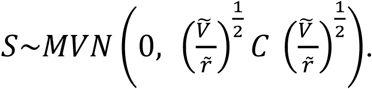

With this formula, Robust SKAT, SKAT-O and burden test can be performed.

### Extension to the joint test of common and rare variants

Our robust method can be extended to the joint test of common and rare variants. Consider the following model

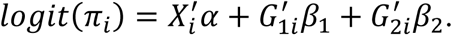

For the individual *i, π*_*i*_ is the disease probability; *X*_*i*_ is the vector containing all the covariates, including the intercept; *G*_1*i*_ is the genotype vector of rare variants with length *m*_*r*_; and *G*_2*i*_ is the vector of common variants with length *m*_*c*_. To test the hypothesis of no genetic effects: *H*_0_: *β*_1_ = 0, *β*_2_ = 0, the test statistic *Q*_*Φ*_ can be written as

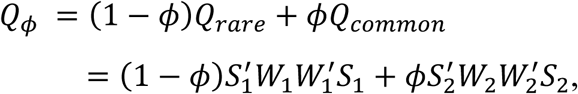

where *S*_1_ and *S*_2_ are the vectors of score statistics f1or rare and com2mon variants respectively, and *W*_1_ and *W*_2_ are diagonal weight matrices for rare and common variants.

Under the null, 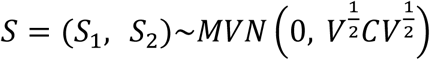. Using the approach described in the previous section, we apply SPA and ER to calibrate variance estimates to perform a robust SKAT method.

### Numerical Simulations

We conducted extensive simulation studies to evaluate the performance of the proposed methods for dichotomized traits. The sequence data of mimicking European ancestry over 200 kb regions were generated using the calibrated coalescent model^29^. We randomly selected regions with lengths of 1 kb and tested for associations in all simulation settings. On average each simulated dataset had 16.33 (SD: 4.05) rare variants when the sample size was 50,000.

We generated data sets with sample size 50,000. We included two covariates for the analysis. The first one followed a Bernoulli distribution with *p* = 0.5 and the other followed the standard normal distribution, corresponding to the gender and normalized age. Four case-control ratios were considered, 1:1, 1:9, 1:49 and 1:99, and the binary phenotypes were simulated from

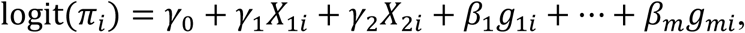

where *β*_1_ = *β*_2_ = *…* = *β*_*m*_ = 0; *γ*_1_ and *γ*_2_ were chosen to let the odds ratio (OR) of *X*_1_ and *X*_2_ equal 1.2 and 1.5 respectively, and *γ*_0_ was chosen based on disease prevalence. Seven different methods were applied to each of the generated dataset. For all variants in the region, we applied the unadjusted and robust joint test of common and rare variants. For rare variant tests (MAF<=0.01), we applied (1) SKAT; (2) robust SKAT; (3) burden test; (4) robust burden test; (5) SKAT-O; (6) robust SKAT-O; and (7) the hybrid method. The hybrid method^16^, developed by Lee, selects a method among ER, Quantile adjusted moment matching (QA) and Moment matching adjustment (MA) based on MAC, and the degree of case-control imbalance. A total of 10^7^ phenotypes were generated, and type I error rates were estimated by the proportion of p-values smaller than the given *α* level divided by given *α*.

For power simulations, 30% of variants were randomly selected as causal variants with the same OR. For each setting, 10,000 data sets were generated, and the power was estimated as the proportion of p-values smaller than the empirical *α* level, which was calculated in the type I error simulation.

### Analysis of whole exome sequencing (WES) data in the UK Biobank

We have analyzed the first tranche of UK Biobank WES data with 49,960 participants^1^. We have downloaded genotype data processed from the Regeneron pipeline. The details of sample selection, variant calling and QC procedures are described elsewhere^1^. We excluded one individual in related pairs (up to second-degree relatives) to identify a set of unrelated individuals. To preserve cases, we first selected a maximal set of unrelated cases, then removed controls that were related to the unrelated cases and kept a maximal set of unrelated controls. Because of the missing values in the phenotypes, the individuals included in the analysis varied across phenotypes. We performed gene-based tests on 45,596 independent European participants in the UK Biobank, whose phenotype data were available.

With a previously published scheme^30^, we defined disease-specific binary phenotypes by combining hospital ICD-9 codes into hierarchical PheCodes, each representing a specific disease group. ICD-10 codes were mapped to PheCodes using a combination of available maps through the Unified Medical Language System (https://www.nlm.nih.gov/research/umls/), manual review and other sources. Study participants were labeled a PheCode if they had one or more of the PheCode-specific ICD codes. Cases were defined as all study participants with the PheCode of interest and controls were all study participants without the PheCode of interest or any related PheCodes. Gender checks were performed, so PheCodes specific for one gender could not be assigned to the other gender by mistake^15^.

There were 791 binary phenotypes with at least 50 cases based on PheCodes, in which 551 phenotypes had case-control ratios smaller than 1:99. Because our robust methods would cause a certain inflation for extremely unbalanced case-control ratios (Supplementary Table 1), we did matching on these 551 traits using the first 4 genotype principal components in which for each case we found the closest controls in Euclidean distance to make the case-control ratio be 1:99. We focused on the rare variants (MAF<=0.01) of the nonsynonymous and splicing variants in the exon and neighboring regions. A total of 18,360 genes were used for the analysis. The number of variants in genes ranged from 2 to 7,439 with a highly skewed distribution (Supplementary Figure 6). Six methods discussed in the simulation study, unadjusted and robust version of SKAT, burden and SKAT-O methods, were applied to the data. Age, gender and first four principal components were used as covariates to adjust for population stratification.

## Code and Data Availability

The proposed robust methods are implemented as an open-source R package available at https://github.com/leeshawn/SKAT/tree/Sparse_Version.

The GWAS results for 791 binary phenotypes with the PheCodes constructed based on ICD codes in UK Biobank using robust SKAT-O are available at http://ukb-50kexome.leelabsg.org, which consists of gene-based Manhattan plots, single variant plots for each gene-phenotype association as well as the PheWAS plots for every gene.

## URLs

Robust gene-based test, https://github.com/leeshawn/SKAT/tree/Sparse_Version. SKAT (version 1.3.2.1), https://cran.r-project.org/web/packages/SKAT

UK-Biobank, https://www.ukbiobank.ac.uk/

UK-Biobank analysis results (gene-based test for binary phenome), http://ukb-50kexome.leelabsg.org/

## Acknowledgements

This research has been conducted using the UK Biobank Resource under application number 45227. SL, ZZ and WB were supported by NIH R01 HG008773.

## Author Contributions

Z.Z. and S.L. designed experiments. Z.Z. and S.L. performed experiments. Z.Z. implemented the software. Z.Z., W.B., W.Z. and L.G.F. analyzed UK Biobank data. P.V. developed the PheWEB like visual server. Z.Z. and S.L. wrote the manuscript.

