## Supplementary Material for "UK-Biobank Whole Exome Sequence Binary Phenome Analysis with Robust Region-based Rare Variant Test"

Supplementary Table 1. Simulation studies of type I error rate of different methods of testing an association with dichotomous traits at stringent $\alpha$ levels $\alpha = 10^{-2}, 10^{-4}$ and $2.5 \times 10^{-6}$.

| | | SKAT | Robust SKAT without additional adjustment | Robust SKAT | Burden | Robust burden without additional adjustment | Robust burden | SKAT-O | Robust SKAT-O without additional adjustment | Robust SKAT-O | Hybrid SKAT-O |
|---|---|---|---|---|---|---|---|---|---|---|---|
| $10^{-2}$ | 1:1 | 0.99 | 0.99 | 0.99 | 1.00 | 1.00 | 1.00 | 1.11 | 1.11 | 1.11 | 1.09 |
| | 1:9 | 1.01 | 1.01 | 1.01 | 0.99 | 0.99 | 1.00 | 1.13 | 1.13 | 1.13 | 1.09 |
| | 1:49 | 1.44 | 1.24 | 1.22 | 1.02 | 0.95 | 0.95 | 1.44 | 1.24 | 1.23 | 1.27 |
| | 1:99 | 1.92 | 1.44 | 1.41 | 1.07 | 0.92 | 0.91 | 1.82 | 1.36 | 1.33 | 1.53 |
| | 1:199 | 2.74 | 1.76 | 1.71 | 1.19 | 0.91 | 0.88 | 2.47 | 1.56 | 1.52 | 2.00 |
| | 1:399 | 3.99 | 2.22 | 2.14 | 1.44 | 0.94 | 0.89 | 3.47 | 1.88 | 1.82 | 2.75 |
| $10^{-4}$ | 1:1 | 0.99 | 0.98 | 1.03 | 1.02 | 1.03 | 1.00 | 1.27 | 1.28 | 1.32 | 1.27 |
| | 1:9 | 1.39 | 1.11 | 1.14 | 1.12 | 1.08 | 0.99 | 1.65 | 1.43 | 1.40 | 1.52 |
| | 1:49 | 6.31 | 1.79 | 1.65 | 2.43 | 1.56 | 0.97 | 6.16 | 2.33 | 1.79 | 4.54 |
| | 1:99 | 13.48 | 2.42 | 2.13 | 3.95 | 2.05 | 1.02 | 12.77 | 3.23 | 2.17 | 8.89 |
| | 1:199 | 28.84 | 3.42 | 2.89 | 6.79 | 2.54 | 1.01 | 26.55 | 4.38 | 2.72 | 17.95 |
| | 1:399 | 61.28 | 4.86 | 3.79 | 11.71 | 2.84 | 0.95 | 55.65 | 5.75 | 3.37 | 36.76 |
| $2.5 \times 10^{-6}$ | 1:1 | 1.24 | 1.03 | 1.54 | 1.11 | 0.94 | 1.03 | 1.38 | 1.19 | 1.38 | 1.40 |
| | 1:9 | 2.47 | 1.19 | 1.45 | 1.29 | 1.41 | 0.77 | 2.51 | 1.61 | 1.49 | 2.23 |
| | 1:49 | 28.27 | 1.80 | 1.91 | 6.88 | 2.91 | 1.06 | 23.70 | 3.24 | 1.98 | 16.69 |
| | 1:99 | 89.53 | 2.67 | 1.81 | 16.34 | 3.80 | 0.90 | 71.32 | 4.87 | 1.60 | 42.59 |
| | 1:199 | 262.60 | 4.62 | 3.16 | 39.57 | 5.99 | 1.17 | 211.90 | 7.93 | 2.80 | 126.26 |
| | 1:399 | 715.05 | 6.70 | 3.90 | 86.87 | 6.65 | 0.80 | 577.70 | 10.70 | 2.83 | 347.10 |

Supplementary Table 2. Simulation studies of type I error rates of different methods of testing an association between all variants, including both common and rare variants, and dichotomous traits at stringent $\alpha$ levels $\alpha = 10^{-2}, 10^{-4}$ and $2.5 \times 10^{-6}$.

| | | SKAT-CommonRare | Robust SKAT-CommonRare without additional adjustment | Robust SKAT-CommonRare |
|---|---|---|---|---|
| $10^{-2}$ | 1:1 | 1.00 | 0.99 | 0.99 |
| | 1:9 | 1.00 | 1.01 | 1.01 |
| | 1:49 | 1.11 | 1.22 | 1.22 |
| | 1:99 | 1.26 | 1.42 | 1.41 |
| | 1:199 | 1.52 | 1.71 | 1.72 |
| | 1:399 | 1.97 | 2.15 | 2.15 |
| $10^{-4}$ | 1:1 | 0.98 | 0.98 | 1.04 |
| | 1:9 | 1.09 | 1.11 | 1.13 |
| | 1:49 | 2.42 | 1.73 | 1.69 |
| | 1:99 | 4.20 | 2.26 | 2.21 |
| | 1:199 | 7.84 | 3.02 | 3.03 |
| | 1:399 | 16.02 | 4.01 | 4.04 |
| $2.5\times10^{-6}$ | 1:1 | 0.94 | 1.05 | 1.66 |
| | 1:9 | 1.53 | 1.17 | 1.45 |
| | 1:49 | 7.48 | 1.57 | 1.94 |
| | 1:99 | 18.54 | 2.50 | 1.93 |
| | 1:199 | 50.43 | 3.87 | 3.37 |
| | 1:399 | 140.09 | 4.47 | 4.46 |

Supplementary Table 3. Gene-phenotype associations detected by Robust SKAT-O across 791 phenotypes at $\alpha=2.5\times10^{-6}$ (Number of associations=117)

| Phenotype | Gene Name | Case | Control | N SNP | Total minor counts for cases | Total minor counts for controls | Robust SKAT | Robust burden | Robust SKAT-O |
|---|---|---|---|---|---|---|---|---|---|
| Myeloproliferative disease | *JAK2* | 94 | 9306 | 68 | 26.0 | 435.0 | 1.35E-30 | 1.94E-13 | 8.92E-30 |
| Cervical intraepithelial neoplasia [CIN] [Cervical dysplasia] | *NAGS* | 309 | 21875 | 81 | 21.0 | 340.0 | 8.63E-05 | 8.94E-09 | 4.71E-09 |
| Large cell lymphoma | *TNC* | 56 | 5544 | 135 | 14.0 | 490.8 | 7.49E-08 | 4.03E-04 | 6.10E-09 |
| Cancer of prostate | *HOXB13* | 741 | 18940 | 37 | 18.0 | 153.6 | 6.90E-08 | 2.15E-05 | 3.00E-08 |
| Spondylosis and allied disorders | *MAP3K7CL* | 849 | 43787 | 58 | 14.0 | 153.0 | 1.49E-07 | 7.58E-06 | 4.85E-08 |
| Congenital coagulation defects | *F11* | 76 | 7524 | 39 | 8.0 | 85.0 | 1.97E-06 | 2.12E-06 | 6.13E-08 |
| Peptic ulcer (excl. esophageal) | *LMNB2* | 773 | 44818 | 171 | 24.0 | 501.0 | 5.39E-07 | 5.39E-06 | 6.65E-08 |
| Menopausal and postmenopausal disorders | *NFE2L3* | 1345 | 21226 | 172 | 144.0 | 1369.0 | 3.79E-06 | 2.17E-07 | 6.93E-08 |
| Other aneurysm | *P3H1* | 164 | 16236 | 112 | 17.0 | 498.0 | 3.49E-07 | 1.12E-05 | 7.13E-08 |
| Congenital anomalies of great vessels | *SLC46A1* | 134 | 13266 | 29 | 11.0 | 256.0 | 7.05E-08 | 4.29E-05 | 7.50E-08 |
| Unspecified monoarthritis | *OGG1* | 1728 | 41060 | 122 | 118.0 | 1650.0 | 1.75E-06 | 3.38E-07 | 1.03E-07 |
| Fracture of hand or wrist | *GSDMC* | 382 | 37818 | 110 | 25.0 | 760.0 | 1.73E-05 | 6.64E-07 | 1.07E-07 |
| Generalized convulsive epilepsy | *KRT6A* | 52 | 5148 | 49 | 15.0 | 275.1 | 1.53E-07 | 2.82E-05 | 1.39E-07 |
| Other disorders of male genital organs | *ARIH1* | 325 | 18835 | 36 | 11.1 | 131.5 | 3.42E-07 | 1.20E-05 | 1.50E-07 |
| Heart valve replaced | *RBM25* | 131 | 12969 | 46 | 10.1 | 171.9 | 3.50E-07 | 4.07E-06 | 2.86E-07 |
| Abnormal findings on exam of gastrointestinal tract/ abdominal area | *OR51T1* | 169 | 16731 | 46 | 13.0 | 205.0 | 8.76E-05 | 1.98E-07 | 2.90E-07 |
| Agorophobia, social phobia, and panic disorder | *GPR182* | 78 | 7722 | 34 | 6.0 | 69.0 | 6.22E-07 | 3.65E-05 | 2.98E-07 |
| Other open wound of head and face | *CRACR2B* | 304 | 30096 | 110 | 9.1 | 424.5 | 1.32E-07 | 3.68E-02 | 3.38E-07 |

| | | | | | | | | | |
|---|---|---|---|---|---|---|---|---|---|
| Other and unspecified disc disorder | *ODF2* | 447 | 43796 | 202 | 48.0 | 2299.0 | 5.66E-07 | 5.84E-06 | 4.46E-07 |
| Disorders of penis | *RINL* | 360 | 18836 | 74 | 12.0 | 164.1 | 3.25E-07 | 4.81E-05 | 4.95E-07 |
| Noninflammatory female genital disorders | *STX17* | 1139 | 23792 | 38 | 39.0 | 337.0 | 1.31E-03 | 6.19E-07 | 5.28E-07 |
| Elevated blood pressure reading without diagnosis of hypertension | *SFXN1* | 167 | 16533 | 45 | 14.0 | 277.0 | 3.31E-05 | 5.95E-07 | 5.62E-07 |
| Benign neoplasm of breast | *C8orf33* | 177 | 17523 | 45 | 15.0 | 336.0 | 4.45E-07 | 9.14E-06 | 5.70E-07 |
| Hearing loss | *REEP3* | 372 | 36828 | 42 | 7.0 | 92.0 | 1.21E-06 | 1.96E-05 | 5.87E-07 |
| Dislocation | *MDFI* | 255 | 25245 | 45 | 10.0 | 169.0 | 1.89E-06 | 8.05E-06 | 6.08E-07 |
| Epilepsy, recurrent seizures, convulsions | *MAK* | 497 | 44306 | 117 | 21.0 | 713.0 | 5.59E-07 | 1.22E-04 | 6.92E-07 |
| Scar conditions and fibrosis of skin | *C1orf109* | 265 | 26235 | 53 | 10.0 | 226.0 | 1.04E-06 | 9.61E-05 | 6.92E-07 |
| Disturbance of skin sensation | *CXCR5* | 289 | 28611 | 55 | 7.0 | 147.0 | 8.33E-07 | 4.52E-04 | 7.98E-07 |
| Edema | *SFPQ* | 160 | 15840 | 64 | 7.0 | 196.2 | 5.98E-07 | 8.07E-03 | 8.14E-07 |
| Redundant prepuce and phimosis/BXO | *RINL* | 296 | 18837 | 73 | 9.0 | 164.1 | 3.41E-07 | 6.00E-04 | 8.39E-07 |
| Cardiac congenital anomalies | *SLC46A1* | 207 | 20493 | 44 | 13.0 | 416.0 | 6.85E-07 | 2.11E-04 | 8.66E-07 |
| Cholelithiasis with other cholecystitis | *DDAH2* | 527 | 43762 | 52 | 8.0 | 185.0 | 3.34E-07 | 1.33E-03 | 8.68E-07 |
| Diseases of the oral soft tissues, excluding lesions specific for gingiva and tongue | *ECH1* | 533 | 44851 | 84 | 14.1 | 243.1 | 2.25E-05 | 1.37E-06 | 9.02E-07 |
| Hyposmolality and/or hyponatremia | *DLX3* | 144 | 14256 | 26 | 12.0 | 174.0 | 2.42E-04 | 5.12E-07 | 9.55E-07 |
| Hypertension complicating pregnancy, childbirth, and the puerperium | *CLDN18* | 173 | 17127 | 35 | 8.0 | 268.0 | 4.29E-07 | 1.20E-02 | 9.75E-07 |
| Malignant neoplasm of female breast | *ST6GALNAC6* | 1307 | 22672 | 63 | 24.0 | 176.0 | 1.46E-06 | 1.35E-04 | 1.03E-06 |
| Influenza | *RPL21* | 52 | 5148 | 3 | 3.0 | 1.0 | 8.47E-06 | 1.27E-06 | 1.06E-06 |
| Coagulation defects | *F11* | 125 | 12375 | 57 | 9.0 | 151.0 | 2.03E-06 | 1.98E-05 | 1.07E-06 |
| Intracerebral hemorrhage | *GPR108* | 66 | 6534 | 50 | 20.0 | 381.0 | 2.42E-06 | 2.29E-06 | 1.11E-06 |
| Erythematous conditions | *FAM129C* | 267 | 26433 | 128 | 24.0 | 685.0 | 1.45E-04 | 2.56E-07 | 1.12E-06 |

| | | | | | | | | | |
|---|---|---|---|---|---|---|---|---|---|
| Respiratory abnormalities | *CBR4* | 72 | 7128 | 27 | 12.0 | 196.0 | 4.63E-06 | 4.67E-06 | 1.13E-06 |
| Osteoarthrosis, localized, primary | *MRPS14* | 1040 | 42647 | 49 | 12.0 | 118.0 | 2.90E-06 | 2.64E-05 | 1.15E-06 |
| Open wounds of head; neck; and trunk | *CRACR2B* | 394 | 39006 | 136 | 9.1 | 558.5 | 4.15E-07 | 1.68E-01 | 1.16E-06 |
| Cancer, suspected or other | *PCDHGB6* | 1683 | 41603 | 167 | 138.0 | 2375.0 | 7.00E-07 | 2.14E-04 | 1.17E-06 |
| Infection/inflammation of internal prosthetic device; implant; and graft | *UFSP1* | 236 | 23364 | 39 | 12.0 | 208.0 | 2.51E-05 | 1.26E-06 | 1.18E-06 |
| Regional enteritis | *FN3KRP* | 189 | 18711 | 40 | 19.0 | 495.0 | 1.70E-05 | 1.35E-06 | 1.19E-06 |
| Polyp of corpus uteri | *CD300E* | 1059 | 23305 | 31 | 14.0 | 48.0 | 2.12E-04 | 5.21E-07 | 1.19E-06 |
| Other symptoms/disorders or the urinary system | *OR10G3* | 2126 | 42946 | 65 | 87.3 | 992.1 | 9.50E-06 | 1.48E-06 | 1.21E-06 |
| Infection of the eye | *SERPINF2* | 87 | 8613 | 33 | 5.0 | 91.0 | 9.56E-07 | 1.93E-03 | 1.22E-06 |
| Retinal detachments and defects | *ADRB2* | 349 | 34551 | 64 | 7.0 | 205.0 | 4.65E-07 | 2.94E-03 | 1.24E-06 |
| Other local infections of skin and subcutaneous tissue | *OR4C6* | 144 | 14256 | 41 | 9.0 | 162.0 | 2.34E-06 | 3.70E-05 | 1.26E-06 |
| Type 2 diabetes with neurological manifestations | *SCAF8* | 58 | 5742 | 71 | 10.0 | 209.0 | 1.72E-06 | 8.23E-05 | 1.27E-06 |
| Spondylosis without myelopathy | *RASA4;RASA4B* | 553 | 43796 | 82 | 21.5 | 710.6 | 1.11E-06 | 2.31E-04 | 1.28E-06 |
| Hyperhidrosis | *ITSN2* | 75 | 7425 | 108 | 19.0 | 461.1 | 4.75E-04 | 6.70E-07 | 1.29E-06 |
| Fever of unknown origin | *HOXB5* | 455 | 45133 | 42 | 6.0 | 98.0 | 9.00E-07 | 3.74E-04 | 1.29E-06 |
| Electrolyte imbalance | *DLX3* | 345 | 34155 | 39 | 17.0 | 404.0 | 2.38E-03 | 9.27E-07 | 1.29E-06 |
| Epistaxis or throat hemorrhage | *LIN7B* | 267 | 26433 | 38 | 6.0 | 104.0 | 1.38E-06 | 7.71E-04 | 1.31E-06 |
| Otitis media | *LRRC23* | 171 | 16929 | 48 | 8.0 | 155.0 | 9.75E-07 | 3.16E-04 | 1.32E-06 |
| Malignant neoplasm of uterus | *DTD1* | 126 | 12474 | 15 | 4.0 | 63.0 | 1.08E-06 | 1.81E-03 | 1.32E-06 |
| Heartburn | *USP45* | 189 | 18711 | 98 | 22.0 | 628.1 | 1.95E-04 | 1.14E-06 | 1.34E-06 |
| Obesity | *MLC1* | 1192 | 44381 | 109 | 73.0 | 1434.0 | 4.50E-05 | 4.98E-07 | 1.37E-06 |
| Cough | *IDH3A* | 337 | 33363 | 42 | 30.0 | 1044.0 | 8.77E-06 | 5.60E-07 | 1.38E-06 |
| Acute pancreatitis | *LONRF1* | 190 | 18810 | 79 | 13.0 | 233.2 | 5.08E-04 | 8.52E-07 | 1.41E-06 |

| | | | | | | | | | |
|---|---|---|---|---|---|---|---|---|---|
| Other open wound of head and face | *LCTL* | 304 | 30096 | 98 | 15.0 | 925.0 | 5.05E-07 | 6.49E-02 | 1.46E-06 |
| Other complications of pregnancy NEC | *ZNF324B* | 390 | 24541 | 101 | 13.0 | 316.0 | 8.41E-07 | 7.61E-03 | 1.48E-06 |
| Noninflammatory female genital disorders | *TUBA3D* | 1139 | 23792 | 82 | 39.1 | 335.6 | 7.19E-04 | 1.33E-06 | 1.49E-06 |
| Nonspecific abnormal findings in stool contents | *SFMBT1* | 184 | 18216 | 83 | 9.0 | 164.0 | 4.52E-06 | 3.32E-05 | 1.50E-06 |
| Nasal polyps | *MPO* | 456 | 43262 | 209 | 50.0 | 2293.1 | 2.14E-05 | 3.69E-06 | 1.51E-06 |
| Noninflammatory female genital disorders | *ACAP1* | 1139 | 23792 | 104 | 19.0 | 224.0 | 3.80E-07 | 3.20E-02 | 1.56E-06 |
| Sicca syndrome | *SFSWAP* | 58 | 5742 | 66 | 13.0 | 371.0 | 2.04E-06 | 1.74E-04 | 1.64E-06 |
| Irregular menstrual cycle | *GTPBP2* | 219 | 21243 | 40 | 6.0 | 49.0 | 7.44E-06 | 8.98E-06 | 1.64E-06 |
| Breast cancer | *ST6GALNAC6* | 1423 | 43066 | 107 | 26.0 | 372.0 | 4.04E-06 | 7.89E-05 | 1.65E-06 |
| Alteration of consciousness | *POTEF* | 53 | 5247 | 87 | 15.0 | 431.7 | 3.33E-06 | 7.12E-05 | 1.65E-06 |
| Unspecified diffuse connective tissue disease | *L2HGDH* | 344 | 34056 | 75 | 8.0 | 152.0 | 1.42E-06 | 1.56E-04 | 1.69E-06 |
| Osteoarthritis; localized | *GTPBP2* | 1827 | 42621 | 69 | 11.0 | 100.0 | 5.58E-07 | 4.19E-03 | 1.70E-06 |
| Symptoms involving digestive system | *ANKRD35* | 2001 | 37239 | 234 | 143.0 | 1786.0 | 1.55E-05 | 4.87E-06 | 1.71E-06 |
| Swelling of limb | *SAFB* | 545 | 44950 | 162 | 14.0 | 451.0 | 9.44E-07 | 1.72E-03 | 1.73E-06 |
| Other symptoms | *ALDH1L1* | 146 | 14454 | 120 | 18.0 | 598.0 | 2.35E-06 | 1.59E-04 | 1.76E-06 |
| Rheumatoid arthritis and other inflammatory polyarthropathies | *PNOC* | 582 | 41021 | 35 | 9.0 | 108.0 | 2.01E-06 | 6.07E-05 | 1.77E-06 |
| Cardiac and circulatory congenital anomalies | *SLC46A1* | 219 | 21681 | 46 | 13.0 | 444.0 | 9.50E-07 | 4.59E-04 | 1.80E-06 |
| Intracranial hemorrhage | *OAS1* | 172 | 17028 | 67 | 6.2 | 180.6 | 5.54E-07 | 5.75E-03 | 1.82E-06 |
| Other retinal disorders | *PRPF3* | 379 | 37521 | 64 | 7.0 | 126.0 | 2.35E-06 | 2.30E-04 | 1.86E-06 |
| Cancer, suspected or other | *PCDHGC5* | 1683 | 41603 | 185 | 50.0 | 820.0 | 1.00E-06 | 9.23E-03 | 1.88E-06 |
| Other disorders of gallbladder | *OR13F1* | 141 | 13959 | 30 | 7.0 | 109.0 | 1.97E-06 | 8.61E-05 | 1.91E-06 |
| Phlebitis and thrombophlebitis of lower extremities | *RELL1* | 463 | 41305 | 49 | 10.0 | 221.0 | 1.51E-06 | 3.88E-04 | 1.98E-06 |
| Osteoarthritis; localized | *DNAJC24* | 1827 | 42621 | 20 | 27.0 | 248.0 | 2.35E-06 | 1.50E-05 | 1.98E-06 |

| | | | | | | | | | |
|---|---|---|---|---|---|---|---|---|---|
| Inflammatory bowel disease and other gastroenteritis and colitis | *GNA15* | 476 | 37276 | 78 | 26.0 | 748.0 | 3.34E-06 | 1.12E-05 | 1.99E-06 |
| Chronic renal failure [CKD] | *ERMN* | 227 | 22473 | 43 | 15.0 | 341.0 | 9.99E-06 | 1.71E-06 | 2.00E-06 |
| Genitourinary congenital anomalies | *SLC4A10* | 171 | 16929 | 75 | 9.0 | 173.0 | 4.47E-06 | 5.32E-05 | 2.00E-06 |
| Breast cancer [female] | *ST6GALNAC6* | 1398 | 22668 | 63 | 25.0 | 176.0 | 3.29E-06 | 1.37E-04 | 2.02E-06 |
| Other specified benign mammary dysplasias | *TRAIP* | 82 | 8118 | 38 | 6.0 | 93.0 | 2.20E-06 | 4.01E-04 | 2.04E-06 |
| Vascular insufficiency of intestine | *COL5A3* | 53 | 5247 | 111 | 10.0 | 226.1 | 3.74E-06 | 1.45E-04 | 2.05E-06 |
| Benign neoplasm of colon | *CCDC47* | 2257 | 43090 | 71 | 40.0 | 455.0 | 2.25E-06 | 4.23E-03 | 2.09E-06 |
| Other inflammatory spondylopathies | *TRIM36* | 171 | 16929 | 80 | 8.0 | 250.0 | 7.77E-07 | 3.39E-03 | 2.12E-06 |
| Renal colic | *SLC10A1* | 279 | 27621 | 85 | 11.0 | 264.0 | 4.74E-06 | 7.69E-05 | 2.14E-06 |
| Myeloproliferative disease | *ASXL1* | 94 | 9306 | 147 | 13.0 | 415.0 | 2.05E-06 | 4.45E-04 | 2.17E-06 |
| Scar conditions and fibrosis of skin | *CYP46A1* | 265 | 26235 | 55 | 9.0 | 120.0 | 9.43E-06 | 9.40E-06 | 2.19E-06 |
| Atherosclerosis of the extremities | *DONSON* | 67 | 6633 | 42 | 9.0 | 199.0 | 2.37E-06 | 2.50E-03 | 2.22E-06 |
| Disturbance of skin sensation | *PVALB* | 289 | 28611 | 19 | 6.0 | 27.0 | 1.13E-04 | 1.15E-06 | 2.23E-06 |
| Poisoning by anticonvulsants and anti-Parkinsonism drugs | *ENTPD6* | 60 | 5940 | 51 | 8.0 | 173.0 | 2.37E-06 | 1.88E-04 | 2.25E-06 |
| Alcohol-related disorders | *KLHL10* | 1223 | 42646 | 74 | 15.0 | 215.0 | 1.19E-06 | 1.45E-03 | 2.26E-06 |
| Spondylosis without myelopathy | *MAP3K7CL* | 553 | 43796 | 57 | 9.0 | 153.0 | 1.85E-06 | 2.19E-04 | 2.28E-06 |
| Elevated blood pressure reading without diagnosis of hypertension | *FKBP5* | 167 | 16533 | 43 | 9.0 | 178.0 | 2.45E-06 | 1.02E-04 | 2.29E-06 |
| Other abnormal glucose | *PIEZO2* | 51 | 5049 | 158 | 15.0 | 389.0 | 1.70E-05 | 1.29E-05 | 2.30E-06 |
| Hypotension NOS | *IDO1* | 356 | 35244 | 70 | 10.0 | 153.0 | 8.71E-06 | 2.08E-05 | 2.34E-06 |
| Disorders of fluid, electrolyte, and acid-base balance | *DDX42* | 675 | 44906 | 125 | 38.0 | 1306.0 | 1.80E-06 | 2.93E-04 | 2.34E-06 |
| Pruritus and related conditions | *FAN1* | 86 | 8514 | 93 | 9.0 | 474.0 | 8.75E-07 | 7.39E-02 | 2.35E-06 |
| Cerebral ischemia | *CHTOP* | 341 | 33759 | 44 | 7.0 | 137.0 | 1.90E-06 | 3.24E-04 | 2.37E-06 |
| Psoriatic arthropathy | *OR9Q1* | 72 | 7128 | 23 | 11.0 | 184.0 | 2.35E-05 | 1.13E-06 | 2.37E-06 |

| | | | | | | | | | |
|---|---|---|---|---|---|---|---|---|---|
| Hypothyroidism NOS | *SERHL2* | 1590 | 43626 | 92 | 69.0 | 1030.0 | 2.09E-05 | 5.75E-06 | 2.41E-06 |
| Congestive heart failure (CHF) NOS | *DUSP2* | 185 | 18315 | 50 | 16.0 | 358.0 | 1.84E-03 | 8.72E-07 | 2.42E-06 |
| Overweight, obesity and other hyperalimentation | *MLC1* | 1210 | 44381 | 109 | 73.0 | 1434.0 | 6.53E-05 | 8.76E-07 | 2.43E-06 |
| Sicca syndrome | *ADAMTS20* | 58 | 5742 | 108 | 10.0 | 181.0 | 1.82E-05 | 1.24E-05 | 2.45E-06 |
| Hypothyroidism | *SERHL2* | 1666 | 43622 | 92 | 72.0 | 1030.0 | 2.92E-05 | 4.19E-06 | 2.47E-06 |
| Fracture of radius and ulna | *SVOP* | 508 | 43549 | 63 | 10.0 | 178.0 | 1.39E-05 | 1.05E-05 | 2.47E-06 |
| Uterine/Uterovaginal prolapse | *AKR1A1* | 563 | 23644 | 50 | 19.0 | 206.0 | 3.43E-04 | 1.24E-06 | 2.48E-06 |
| Viral hepatitis | *SLC28A1* | 73 | 7227 | 56 | 6.0 | 142.0 | 1.01E-06 | 2.23E-03 | 2.50E-06 |

Supplementary Table 4. Top 3 single rare-variant signals of associations with p-value < $10^{-7}$ in the UK Biobank WES data.

| Phenotype (Phecode) | Gene Name | Location | MAF | p-value | Annotation | Polyphen | SIFT |
|---|---|---|---|---|---|---|---|
| Myeloproliferative disease (200) | *JAK2* | 9:5073770:G:T | 1.28E-03 | 4.39E-36 | nonsynonymous SNV | probably_damaging | deleterious |
| | | 9:5070022:D:6 | 5.32E-05 | 5.87E-03 | nonframeshift deletion | - | - |
| | | 9:5022213:G:T | 1.06E-04 | 7.16E-03 | nonsynonymous SNV | probably_damaging | deleterious |
| Cervical intraepithelial neoplasia [CIN] [Cervical dysplasia] (180.3) | *NAGS* | 17:44005037:C:A | 3.94E-03 | 4.37E-04 | nonsynonymous SNV | benign | tolerated |
| | | 17:44007442:G:A | 4.51E-04 | 1.22E-03 | nonsynonymous SNV | possibly_damaging | deleterious |
| | | 17:44004856:G:A | 1.80E-04 | 3.43E-03 | nonsynonymous SNV | benign | tolerated_low_confidence |
| Large cell lymphoma (202.24) | *TNC* | 9:115084397:C:T | 1.52E-03 | 1.02E-05 | nonsynonymous SNV | probably_damaging | tolerated |
| | | 9:115087220:C:G | 2.05E-03 | 1.35E-05 | nonsynonymous SNV | probably_damaging | tolerated |
| | | 9:115087037:C:T | 2.68E-04 | 1.00E-02 | nonsynonymous SNV | probably_damaging | tolerated |
| Cancer of prostate (185) | *HOXB13* | 17:48728343:C:T | 2.16E-03 | 5.24E-08 | nonsynonymous SNV | probably_damaging | deleterious |
| | | 17:48728250:G:A | 7.62E-05 | 3.37E-02 | nonsynonymous SNV | benign | deleterious |
| | | 17:48728379:C:A | 1.02E-04 | 7.05E-02 | nonsynonymous SNV | possibly_damaging | tolerated |
| Spondylosis and allied disorders (721) | *MAP3K7CL* | 21:29174807:G:A | 1.23E-04 | 2.11E-07 | nonsynonymous SNV | benign | tolerated |
| | | 21:29092522:T:A | 7.84E-05 | 8.45E-05 | nonsynonymous SNV | benign | deleterious_low_confidence |
| | | 21:29133376:A:G | 1.12E-05 | 6.45E-03 | nonsynonymous SNV | - | - |
| Congenital coagulation defects (286.1) | *F11* | 4:186288589:T:G | 1.32E-04 | 4.52E-05 | nonsynonymous SNV | probably_damaging | deleterious |
| | | 4:186273178:G:A | 1.97E-04 | 1.73E-04 | splicing | LoF: High-confidence | - |
| | | 4:186267139:G:T | 6.58E-05 | 3.85E-03 | nonsynonymous SNV | probably damaging | deleterious |
| Peptic ulcer (excl. esophageal) (531) | *LMNB2* | 19:2434094:G:A | 2.30E-04 | 3.83E-06 | nonsynonymous SNV | probably_damaging | deleterious |
| | | 19:2456926:G:T | 2.19E-05 | 1.47E-04 | nonsynonymous SNV | - | - |
| | | 19:2431845:C:A | 1.10E-05 | 3.45E-03 | nonsynonymous SNV | benign | deleterious |
| Menopausal and postmenopausal disorders (627) | *NFE2L3* | 7:26184630:A:G | 7.93E-03 | 2.72E-05 | nonsynonymous SNV | benign | deleterious |
| | | 7:26185140:G:A | 5.91E-03 | 2.18E-03 | nonsynonymous SNV | benign | tolerated |
| | | 7:26185559:A:G | 3.33E-04 | 6.71E-03 | nonsynonymous SNV | benign | tolerated |
| Other aneurysm (442) | *P3H1* | 1:42766778:C:T | 6.71E-04 | 1.71E-05 | nonsynonymous SNV | probably_damaging | tolerated |
| | | 1:42748243:C:T | 3.35E-04 | 1.21E-04 | nonsynonymous SNV | benign | tolerated |
| | | 1:42758970:C:G | 9.15E-05 | 3.88E-04 | nonsynonymous SNV | probably_damaging | deleterious |
| Congenital anomalies of great vessels (747.13) | *SLC46A1* | 17:28405185:A:T | 1.75E-03 | 1.86E-08 | nonsynonymous SNV | possibly_damaging | deleterious |
| | | 17:28402276:C:T | 3.73E-05 | 1.24E-02 | nonsynonymous SNV | possibly_damaging | deleterious |
| | | 17:28405926:C:G | 5.22E-04 | 7.13E-01 | nonsynonymous SNV | benign | tolerated |

Supplementary Table 5. The most significant nearby variant association signals (± 100 Kbp up and down stream) in the UK-Biobank imputed datasets of 400,000 British samples.

| Phenotype (Phecode) | Gene Name | Location | Ref Allele | Alt Allele | MAF | p-value |
|---|---|---|---|---|---|---|
| Myeloproliferative disease (200) | *JAK2* | chr9:5075628 | C | G | 2.51E-01 | 2.30E-17 |
| Cervical intraepithelial neoplasia [CIN] [Cervical dysplasia] (180.3) | *NAGS* | chr17:44018609 | A | G | 7.69E-05 | 2.12E-03 |
| Large cell lymphoma (202.24) | *TNC* | chr9:114933743 | G | A | 5.04E-04 | 7.51E-05 |
| Cancer of prostate (185) | *HOXB13* | chr17:48733224 | C | T | 3.27E-02 | 1.12E-04 |
| Spondylosis and allied disorders (721) | *MAP3K7CL* | chr21:29029344 | C | T | 2.62E-03 | 2.64E-03 |
| Congenital coagulation defects (286.1) | *F11* | chr4:186258582 | G | C | 4.12E-04 | 4.21E-03 |
| Peptic ulcer (excl. esophageal) (531) | *LMNB2* | chr19:2422611 | C | T | 4.40E-05 | 8.81E-04 |
| Menopausal and postmenopausal disorders (627) | *NFE2L3* | chr7:26175486 | T | A | 3.48E-01 | 2.14E-04 |
| Other aneurysm (442) | *P3H1* | chr1:42751787 | G | A | 9.41E-03 | 5.12E-04 |
| Congenital anomalies of great vessels (747.13) | *SLC46A1* | chr17:28413060 | C | T | 5.98E-03 | 3.71E-04 |

Supplementary Figure 1. Empirical power estimates for the unadjusted and robust version of SKAT and burden test where 30% of variants were causal variants and all causal variants were deleterious. The X-axis represents the genetic effect odds ratio and the Y-axis represents the empirical power. All causal variants had the same odds ratios.

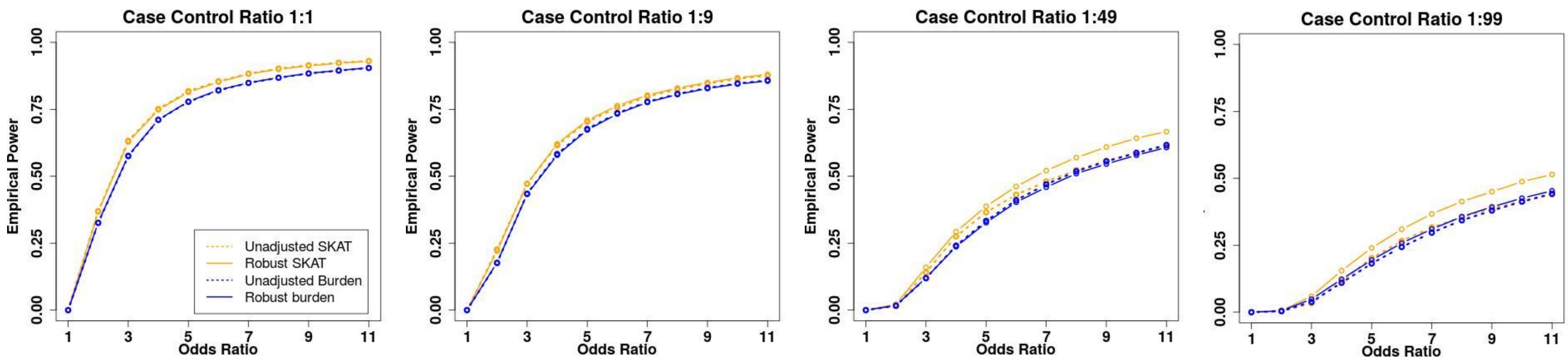

Supplementary Figure 2. Empirical power estimates for the Robust SKAT, burden, SKAT-O with the same number of cases across different case control ratios where 30% of variants were causal variants and all causal variants were deleterious. The X-axis represents the genetic effect odds ratio and the Y-axis represents the empirical power. All causal variants had the same odds ratios.

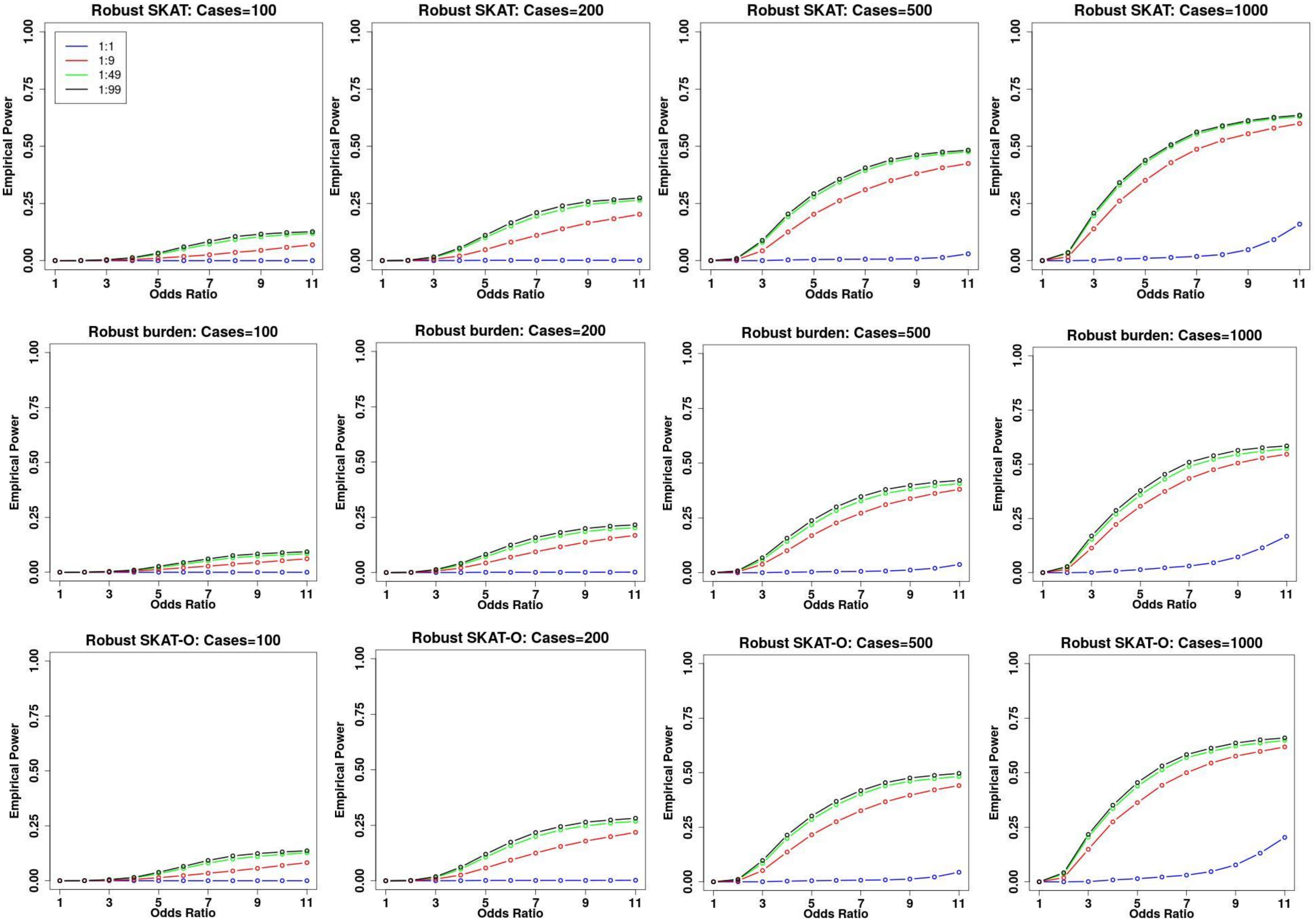

Supplementary Figure 3. Empirical power estimates for robust SKAT, burden and SKAT-O where 30% of variants were causal variants and all causal variants were deleterious. The X-axis represents the genetic effect odds ratio and the Y-axis represents the empirical power. All causal variants had the same odds ratios.

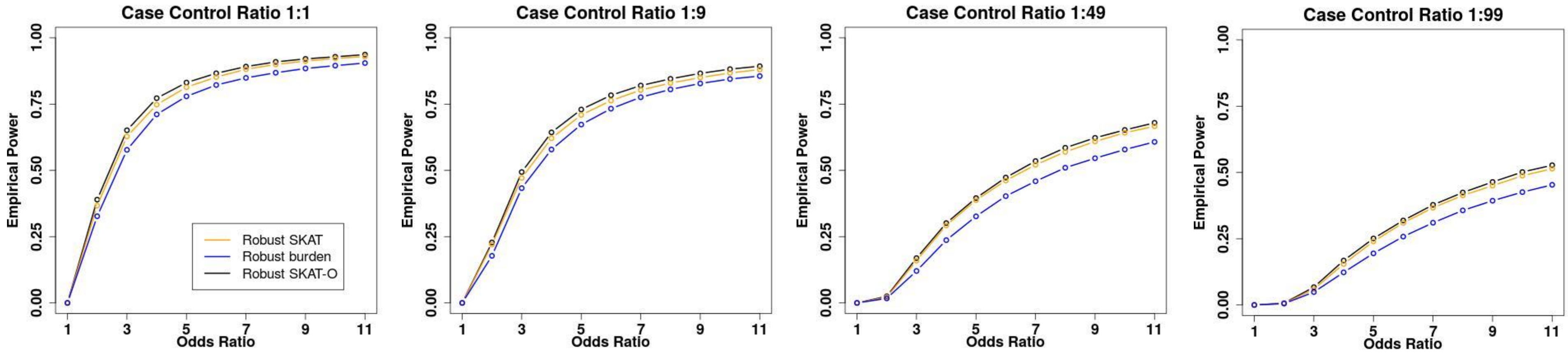

Supplementary Figure 4. P-values of single variants in 10 significant genes. The X-axis represents the position of each single variant in the gene, and the Y-axis represents the negative log10 p-values of single variants. The dashed line represents the cutoff of $5\times10^{-8}$.

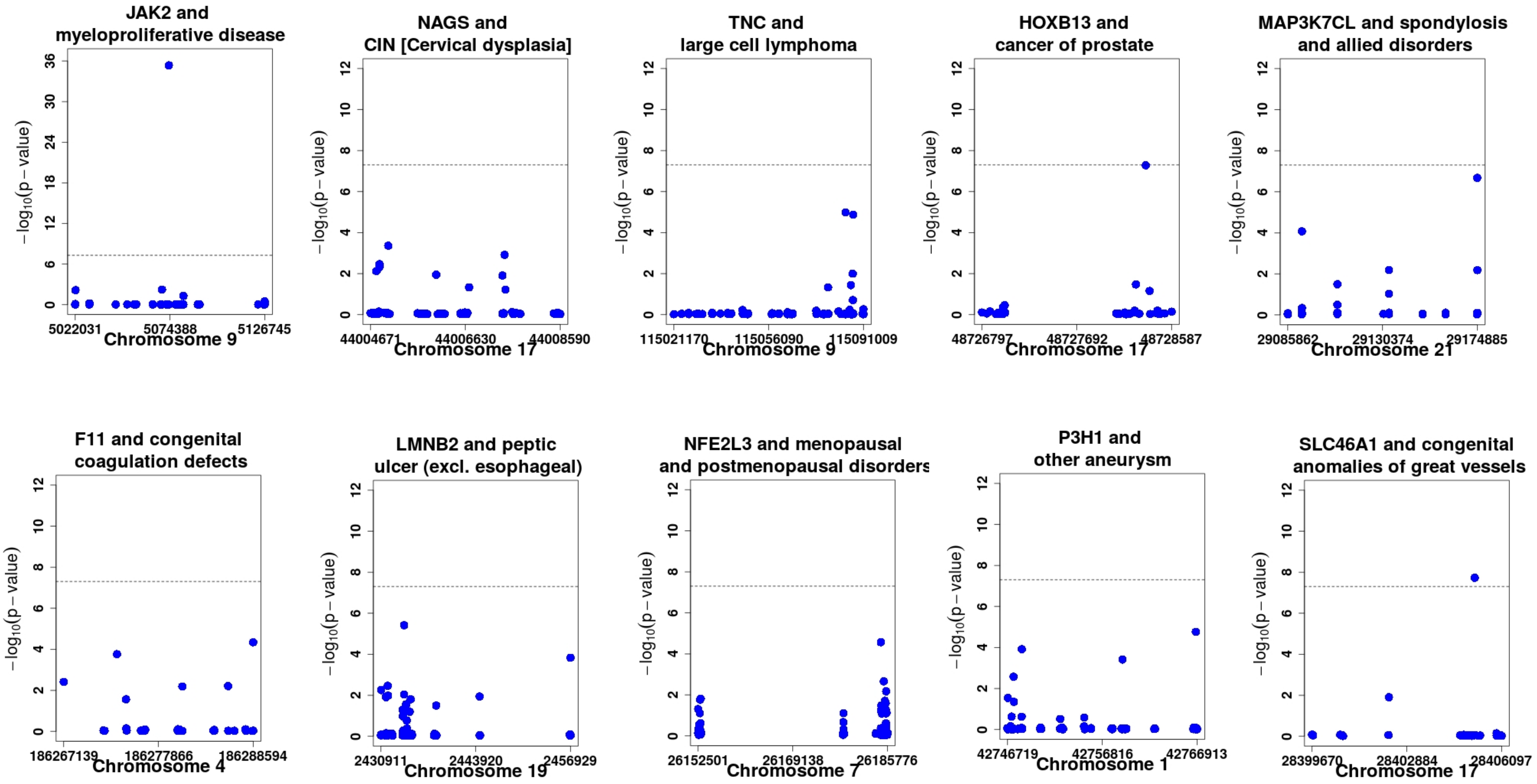

Supplementary Figure 5. QQ Plots of SKAT-O p-values of 10 selected phenotypes. The X-axis represents the expected negative log10 p-values and the Y-axis represents the observed negative log10 p-values of genes.

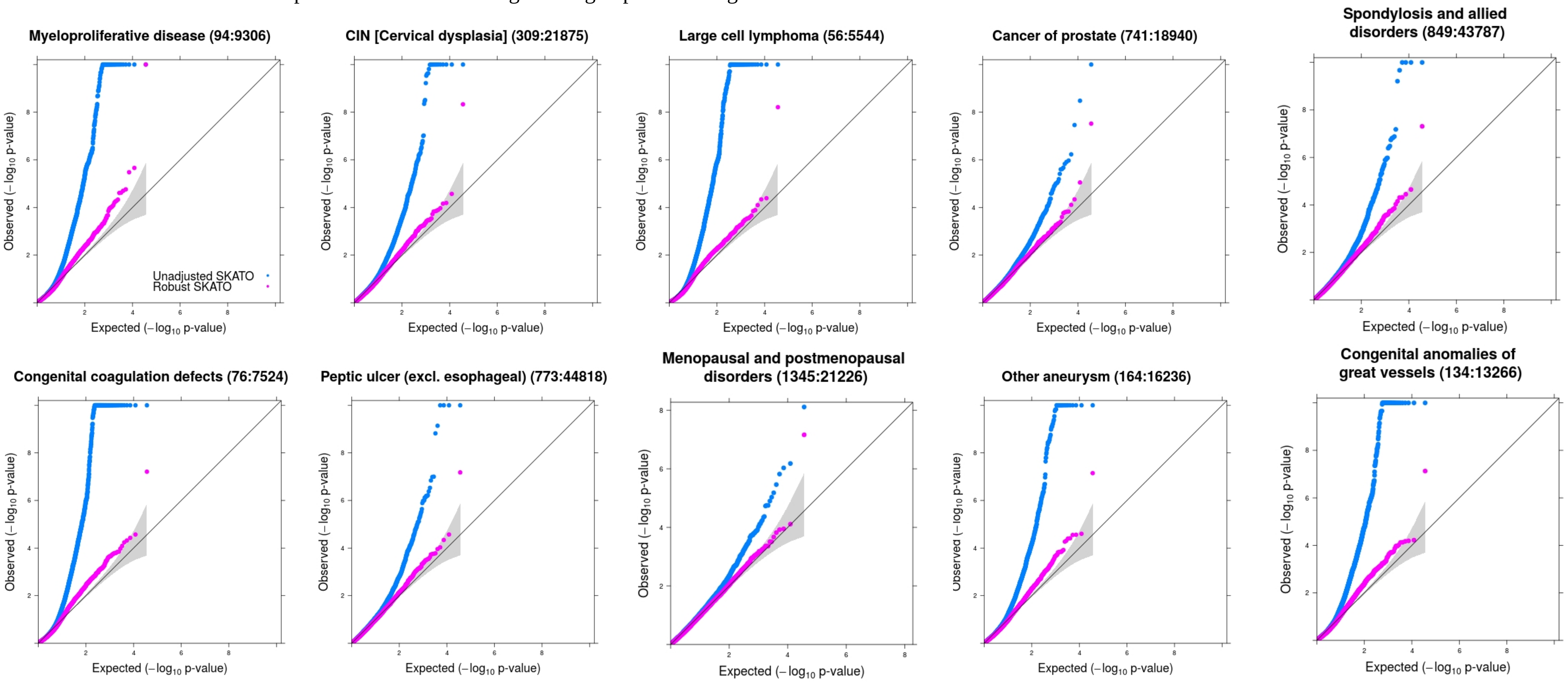

Supplementary Figure 6. The distribution of the number of variants in genes in the UK-Biobank WES data.

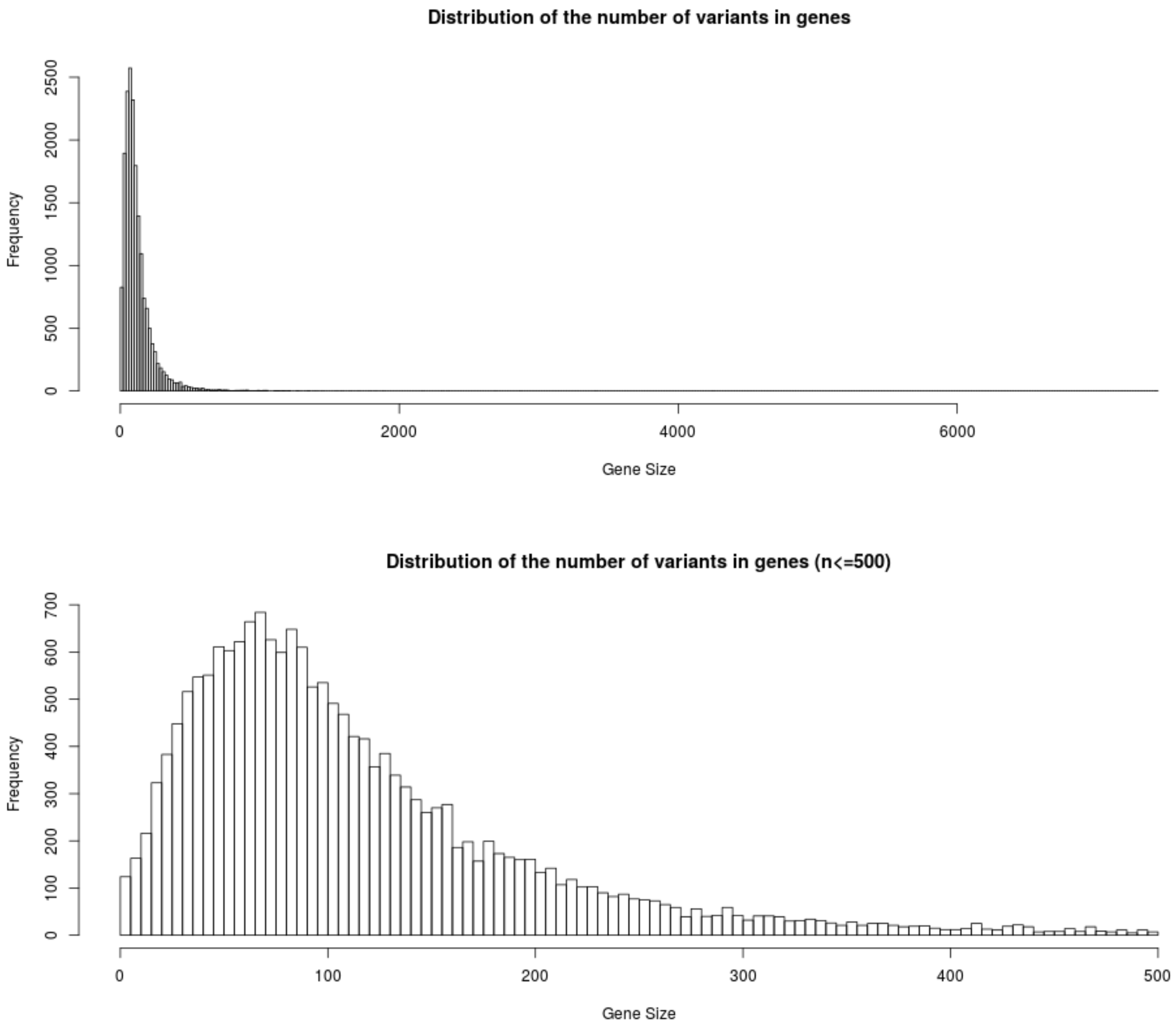
